# Implications of SARS-CoV-2 mutations for genomic RNA structure and host microRNA targeting

**DOI:** 10.1101/2020.05.15.098947

**Authors:** Ali Hosseini Rad SM, Alexander D. McLellan

## Abstract

The SARS-CoV-2 virus is a recently-emerged zoonotic pathogen already well adapted to transmission and replication in humans. Although the mutation rate is limited, recently introduced mutations in SARS-CoV-2 have the potential to alter viral fitness. In addition to amino acid changes, mutations could affect RNA secondary structure critical to viral life cycle, or interfere with sequences targeted by host miRNAs. We have analysed subsets of genomes from SARS-CoV-2 isolates from around the globe and show that several mutations introduce changes in Watson-Crick pairing, with resultant changes in predicted secondary structure. Filtering to targets matching miRNAs expressed in SARS-CoV-2 permissive host cells, we identified twelve separate target sequences in the SARS-CoV-2 genome; eight of these targets have been lost through conserved mutations. A genomic site targeted by the highly abundant miR-197-5p, overexpressed in patients with cardiovascular disease, is lost by a conserved mutation. Our results are compatible with a model that SARS-CoV-2 replication within the human host could be constrained by host miRNA defence. The impact of these and further mutations on secondary structures, miRNA targets or potential splice sites offers a new context in which to view future SARS-CoV-2 evolution, and a potential platform for engineered viral attenuation and antigen presentation.

## Introduction

The SARS-CoV-2 virus has rapidly emerged as a zoonotic pathogen with broad cellular tropism in human, or zoonotic-host cells. Host selection pressure on SARS-CoV-2 virus would be expected to have a major impact on the conservation of mutations that enhance viral fitness. Of these selection pressures, the cellular-based adaptive and innate immune systems will be major constraints to viral evolution. Intracellular detection and anti-viral pathways within infected cells are a critical frontline to control virus replication. The success of the SARS coronaviruses is proposed to be due to their ability to suppress intracellular anti-viral pathways. For example, interference with dsRNA detection and the interferon response is enabled through the activity of several non-structural proteins (Nsp). In addition, the sequestration of genomic viral RNA into double membrane vesicles, and dsRNA cleavage by Nsp15, is inferred from the closely related SARS-CoV-1 virus, and likely acts to prevent intracellular detection of the virus [1]. In addition to encoded mechanisms of immune avoidance, the paucity of CpG runs in SARS-CoV-2 genome with unexpectedly low GC-content at codon position-three points to major selection pressure being placed on structural features of the genome [2].

As a recently emerged zoonotic pathogen, it might be expected that bat-adaptations will not be optimal for infection and replication in human cells. However, extensive mutation and strain-radiation has not yet been observed [3]. The low mutation rate in SARS-CoV-2 is reduced by the activity of the 3’ - 5’ exonuclease Nsp14 in the RNA-dependent RNA polymerase (RdRp) complex. An alternative possibility is that the observed mutation rate is lower than the actual mutation rate, but that deleterious mutations have already been lost through natural selection. The short time frame of SARS-CoV-2 evolution, coupled to a low mutation rate is consistent with a founder effect for geographical bias in mutation patterns [3].

A common primary focus of mutational analysis of emerging viruses is the alteration in amino acid sequence of viral proteins that may provide enhanced or new functions for virus replication, immune avoidance, or spread. However, nucleic acid secondary structure and sub-translational events have a critical impact on genome replication [4], virus maturation and genome packaging [5]. In addition, the RNA secondary structures of SARS-CoV-2 genes have been proposed to be druggable targets [6-8]. Because little is known of the influence of SARS-CoV-2 mutations on the RNA secondary structure, and its possible implications for inhibition by host miRNA, we have modeled the impact of common mutations of the structure and susceptibility of the SARS-CoV-2 genome to interference from host miRNA.

The incident presence of host miRNA targets within the SARS-CoV-2 genome may be pivotal for the action of host selection pressures to further shape further viral evolution. Viruses not only alter host miRNA expression, but may also produce miRNAs to promote their infectivity [9-12]. On the other hand, the host targets viral transcripts for inhibition of translation, or mRNA destruction, through an miRNA-mediated defence system. Since miRNA are extremely divergent between species, it would be expected that bat-adapted SARS-CoV-2 will undergo selection pressure derived from human miRNA interference [9-11,13,14]. While perfect matches of miRNA to target viral sequences results in miRNA-induced silencing complex (miRISC)-mediate destruction of viral RNA, imperfect matches interfere with translation [15].

A growing body of evidence suggests that human miRNAs act as a critical host defense against coronaviruses. An interaction between human coronavirus OC43 nucleocapsid and miR-9 can enhance the type I interferon response necessary to clear viral infection [16]. Several host miRNAs (miR-574-5p, -14, -17, -9, -223 and -148a) bind to SARS-CoV encoded transcripts such as S, E, M, N and ORF1a [17,18]. On the other hand, SARS-CoV escapes from miRNA-mediated defence through the manipulation of host miRNA machinery [17,18]. Additionally, SARS-CoV and SARS-CoV-2 express short RNAs that resemble miRNAs and could impact upon host house-keeping or immune defence processes [19-21]. More recently, several studies have proposed that host miRNAs bind SARS-CoV-2 transcripts [19,21,22]. However, host miRNAs inhibition of viral replication is relevant only if the identified miRNAs are expressed in target host cells.

Both DNA viruses, and ‘cytoplasmically-confined’ RNA viruses, use the host RNA splicing-machinery to generate new viral transcripts, or to modify the host transcriptome in favor of their own replication [23-27]. It has been suggested that the fused leader sequence in 5′ end of the mouse hepatitis virus (betacoronavirus) mRNAs is result of a non-canonical splicing process [28]. Moreover, deep RNA sequencing has identified several unknown SARS-CoV-2 viral RNAs, possibly the result of non-canonical splicing events [29]. Therefore, our study has additionally identified and mapped mutations to predicted mRNA splice sites within the SARS-CoV-2 genome.

No selective advantage of the identified sequence alterations in SARS-CoV-2 should be inferred by their inclusion here. However, the potential of these mutations to impact upon RNA structure and miRNA recognition provides a basis for ongoing monitoring of viral evolution at these sites in the SARS-CoV-2 genome.

The interplay of viral genome sequences and host miRNA might be veiwed as academic and refractory for translation into usable clinical outcomes. However, the inclusion of host miRNA binding sites into the ORF of conserved viral regions essential for the viral life cycle is a feasible mechanism for enhancing the attenuation of live vaccines [30-32], or antigen availability of epiptopes for the adaptive immune response [33-36].

## Methods

SARS-CoV-2 virus reference sequence was downloaded from NCBI (NC_045512.2) along with 55 sequences up to May 1^st^, 2020 from NCBI or GASID databases. We tried to include wide range of countries with available sequences up to May 1^st^. In case of USA, 15 sequences from different states was included. Clustal Omega and Geneious alignment tools were used to perform multiple sequence alignment. Mutations with occurrence in multiple sequences originated from different countries were categorized as conserved mutation.

Potential splice donor / acceptor splice sites, exon splicing enhancer (ESE), exon splicing silencer (ESS), intron splicing enhancer (ISE) and intron splicing silencer (ISS) motifs were predicted using RegRNA2 [37], HSF [38] and NIPU [39,40] tools.

We used well-accepted methods to predict the RNA secondary structure in both wild type and mutated sequences. Minimum free energy (MFE) structures [41] and centroid structures [42] were calculated by RNAfold program to predict RNA secondary structures. To evaluate the impact of mutations on RNA secondary setructure and base-pair probability, we utilized RNAfold, RNAalifold [43], MutaRNA [40,44] and RNAsnp [45] programs.

For identifying potential miRNA binding sites, the SARS-COV-2 genome was screened with RegRNA2 and miRDB [46]. We excluded miRNAs that are not expressed in SARS-CoV-2 target cells such as lung, esophagus, kidney and small intestine [47,48]. The expression level of miRNAs in target cells were determined by TissueAtlas [49], IMOTA [50], or using published data. The impact of mutations on miRNAs binding were visualized by RegRNA2, miRDB, IntaRNA [51] and CopomuS [40]. We used IntaRNA to illustrate miRNA binding to its target.

## Results

### Identification of SARS-Cov-2 conserved mutations

A total of 55 SARS-CoV-2 patient isolate sequences were collected from NCBI and GISAID databases were aligned against SARS-CoV-2 reference sequence (NC_045512.2). The mutations present in multiple sequences and in at least in three different countries were categorized as ‘conserved mutations’ [52]. In the line with previous reports, we confirmed the occurrence of conserved mutations at positions 1397 (Nsp2), 3037 (Nsp3), 8782 (Nsp4), 11083 (Nsp6), 14408 (Nsp12), 23403 (S), 26143 (ORF3a) and 28144/ 28881 (N) [52-56]. We also considered nine additional conserved mutations with occurrence in multiple countries (Table 1 & S1). Most of these mutations are substitutions of C/G to U. The high A/U content (U= 32.1%/ A= 29.9%/ G= 19.6%/ C=18.4%) and enrichment of codons in in pyrimidines is likely due to APOBEC editing of viral RNA and the fact that Nsp14 (proof-reading) does not remove U (the product of cytosine deamination) [57]. Two mutations at 241 and 29742, are in the 5’ and 3’ untranslated regions (UTRs). Seven mutations are silent point mutations, including 313, 14805, 17247 and 28686, while the others result in amino acid changes (Table 1). None of these amino acid changes are conservative substitutions. Interestingly, the C27964U (S24L in ORF8) exists only in 97 USA sequences, with the earliest isolated in March 9^th^ (MT325581.1), after USA underwent lockdown [58] (Figure S1).

**Table 1.** Conserved mutations in SARS-CoV-2 genome.

| **Gene** | **Mutation** | **Amino acid change** |
| --- | --- | --- |
| **5' UTR** | C to U - nt241 | - |
| **Nsp1** | C to U - nt313 | No (L16) |
| **Nsp2** | C to U - nt1059 | T85I |
|  | G to A - nt1397 | V198I |
|  | Deletion 1606-1609 | D268 deletion |
| **Nsp3** | C to U - nt3037 | No (F106) |
| **Nsp4** | C to U - nt8782 | No (S76) |
| **Nsp6** | G to U - nt11083 | L37F |
| **Nsp12** | C to U - nt14408 | P232L |
|  | C to U - nt14805 | No (Y455) |
| **Nsp13** | U to C - nt17247 | No (R337) |
| **S** | C to A - 22277 | Q239K |
|  | A to G - nt23403 | D614G |
|  | C to U - nt24034 | No (N824) |
| **ORF3a** | G to U - nt25563 | Q57H |
|  | G to U - nt26144 | G251V |
| **M** | C to U - nt27406 | T175M |
| **ORF8** | C to U - nt27964 | S24L |
|  | U to C - nt28144 | L84S |
| **N** | C to U - nt28311 | P13L |
|  | U to C - nt28688 | No (L139) |
|  | GGG to AAC - nt28881-28884 | R203K and G204R |
| **3' UTR** | G to U - nt29742 | - |

### RNA secondary structure

Among all the mutations, only two mutations were predicted to have an impact on secondary structure of viral RNA. First, a conserved mutation 1059 in Nsp2 changed the secondary structure of Nsp2 dramatically (Figure 1A). This mutation also increased the Watson-Crick base-pair probability – predicted to result in more stable RNA secondary structure (Figure 1B, C, D & E). This mutation had no effect on RNA accessibility which is a consideration for RNA-RNA and RNA-protein interactions (Figure 1F).

**Figure 1.**
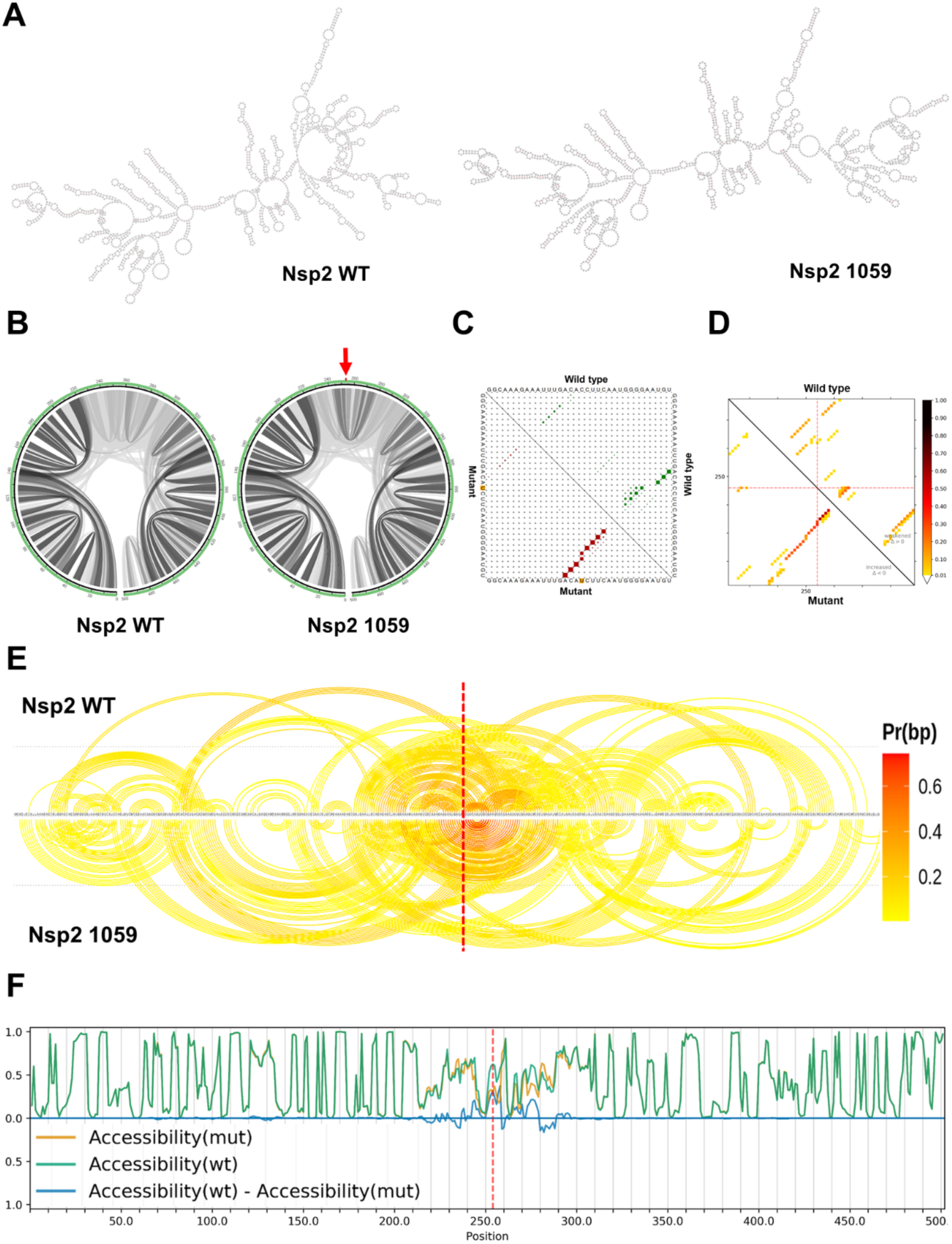
The impact of C1059U mutation on Nsp2 RNA. **(A)** RNA secondary structures of Nsp2 wild type (MFE structure: -571.60 kcal/mol - centroid structure: -491.80 kcal/mol) and 1059 mutation (MFE structure: -572.80 kcal/mol - centroid structure: -512.10 kcal/mol) using RNAfold tool, shows that the 1059 mutation changes the secondary structure of Nsp2 RNA. **(B)** The base pair probabilities by circular plots with higher base pairing potential is reflected in darker hues of gray lines and the mutated position highlighted by red arrow (MutaRNA). **(C)** The upper and lower triangle of the matrix represents the base pair probabilities of wild-type and mutant sequences, respectively, as visualized by RNAsnp (p value = 0.2617, p value < 0.2 is significant). **(D)** The dot plot provides the absolute differences of the base pairing probabilities of mutated vs. wildtype RNA. The upper right part of (top-left) visualizes weakening of the base pairing potential, while the lower left reports base pairs with increased probability within the mutant. Again, darker hues reflect higher absolute changes. The mutated position is highlighted by red dotted lines. **(E)** Arc plot representations of the base pairing probabilities are analogous to the circular plots, but use a heatmap-like arc coloring. The highest probabilities are dark-red while low probable base pairs are depicted in yellow. The 1059 position is marked by a red line. **(F)** The accessibility profiles of wild type (green line) and mutation (yellow line) and their differences provide an assessment of the mutation effect on the RNA single-strandedness, which may relate to its interaction potential with other RNAs or proteins. Accessibility is measured in terms of local single-position unpaired probabilities [59].

Mutation 29742 occurs in a conserved region within 3′ UTR known as the coronavirus 3’ stem-loop II-like motif (s2m). This mutation has significant effect on secondary structure of the 3′ UTR (Figure 2A & B). It is well-known that s2m present in most Coronaviruses and plays a vital role in viral replication and invasion [60-62]. Mutations in this region has been shown to increase the stability of 3′ UTR and its interaction with 5′ UTR [61]. G2974U mutation in s2m causes deletion of weak base-pairing and enhances the appearance of stronger base-pairing, which result in the loss of a weak loop (Figure 2C, D, E & F). The G29742U mutation increases the RNA accessibility in s2m (Figure 2G), whilst removing a c-Myc binding site (GCC AC<u>G</u> CGG A). Several SARS-CoV-2 encoded genes bind to the host proteins involved in biological processes, such as protein trafficking, translation, transcription and ubiquitination regulation [63,64]. In addition, s2m interacts with viral and host proteins such as the polypyrimidine tract-binding protein (PTB), to regulate viral replication and transcription [61,62].

**Figure 2.**
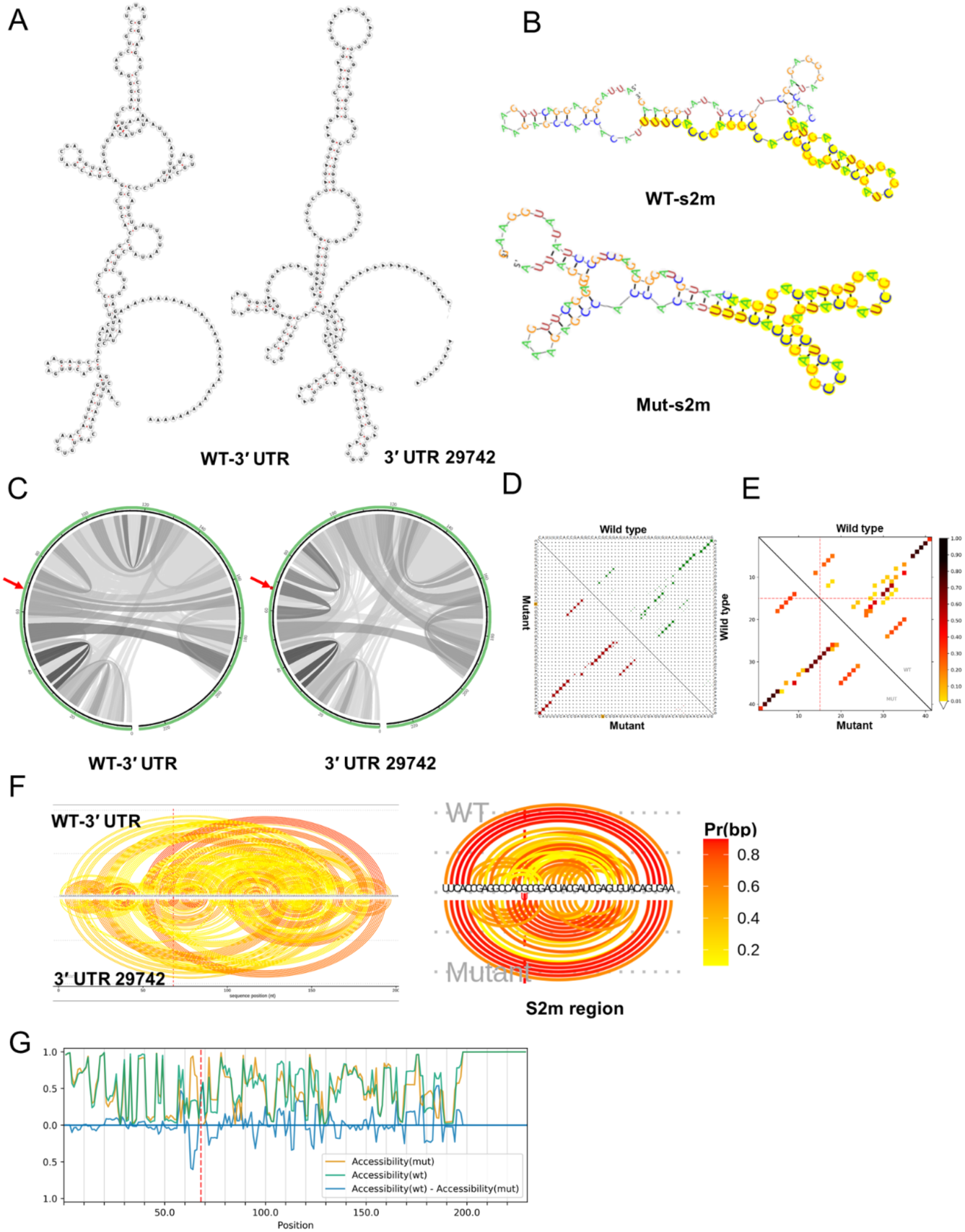
The impact of G29742U mutation on 3′ UTR. **(A)** The RNA secondary structures of wild type 3′ UTR (MFE structure: -36.90 kcal/mol - centroid structure: -30.50 kcal/mol) and 29742 mutation (MFE structure: -40.30 kcal/mol - centroid structure: -30.30 kcal/mol) using RNAfold tool, show that the 29742 mutation changes the secondary structure of 3′ UTR RNA. **(B)** RNA secondary structures of wild type s2m (MFE structure: -6.10 kcal/mol - centroid structure: -0.47 kcal/mol) and 29742 mutation (MFE structure: -11.70 kcal/mol - centroid structure: -11.40 kcal/mol) using RegRNA tool (RegRNA uses RNAfold to creat secondary structures), shows that 29742 mutation changes the secondary structure of s2m region. **(C)** The base pair probabilities by circular plots with higher base pairing potential reflected in darker hues of graduated gray lines. The mutated position is highlighted by a red arrow (MutaRNA). **(D)** The upper and lower triangle of the matrix represents the base pair probabilities of wild-type and mutant sequences, respectively visualized by RNAsnp (p value = 0.6204, p value < 0.2 is significant). **(E)** The dot plot provides the absolute differences of the base pairing probabilities of mutated vs. wild type RNA. Darker hues reflect higher absolute changes. The mutated position is highlighted by red dotted lines. **(F)** Arc plot representations of the base pairing probabilities are analogous to the circular plots, but use a heatmap-like arc coloring. That is highest probabilities are dark red, while low probable base pairs are depicted in yellow. The 29742 position is marked by a red line. **(F)** The accessibility profiles of wild type (green line) and mutation (yellow line) and their differences provide an assessment of the mutation’s effect on the RNA single-strandedness. Accessibility is measured in terms of local single-position unpaired probabilities [59]. The mutated position is highlighted by a red line.

Collectively, these results suggest that 1059 and 29742 yield more stable RNA structures. However, the relationship of changes in RNA secondary structure of Nsp2 and 3′ UTR to viral replication or infectivity must be tested in adequate experimental assays.

### Potential interaction of SARS-CoV-2 transcripts and human miRNAs

In addition to previous studies, we have identified several human miRNAs with potential binding sites across SARS-CoV-2 genome. We filtered our considered miRNA to those with documented expression in SARS-CoV-2 target cells (Figure 3 & S2-11) and additionally focused on miRNAs that have been reported as components of the anti-viral miRNA-mediated defence system. We hypothesised that some mutations may represent escape mutations to avoid miRNA-mediated defence. As shown in Figure 4, ten mutations occur in miRNA binding site and abolish the binding of miRNAs to their targets. Only one mutation (C15293U) inside Nsp12 did not destroy the miR-1273d binding site (GISAID: EPI_ISL_41922).

**Figure 3.**
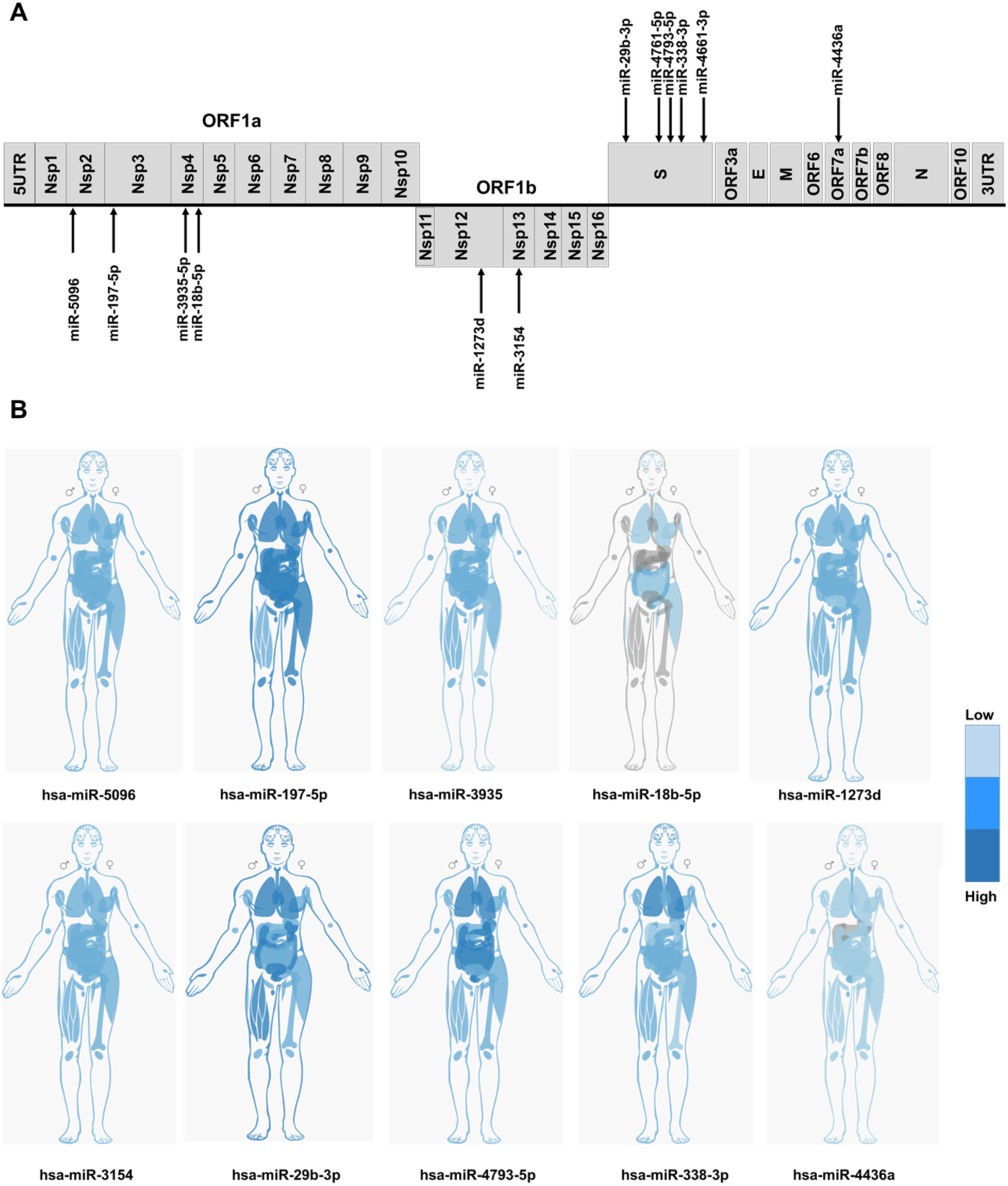
**(A)** Identification of host miRNA targeting different regions of SARS-CoV-2 genome. **(B**) The expression level of candidate miRNA in different human tissues. Data obtained from IMOTA database. Darker blue indicates the higher expression. Grey colour shows zero expression in those tissues. Also, the graph presentation of miRNA expression in different human tissues obtained from TissueAtlas database is available in supplementary data.

**Figure 4.**
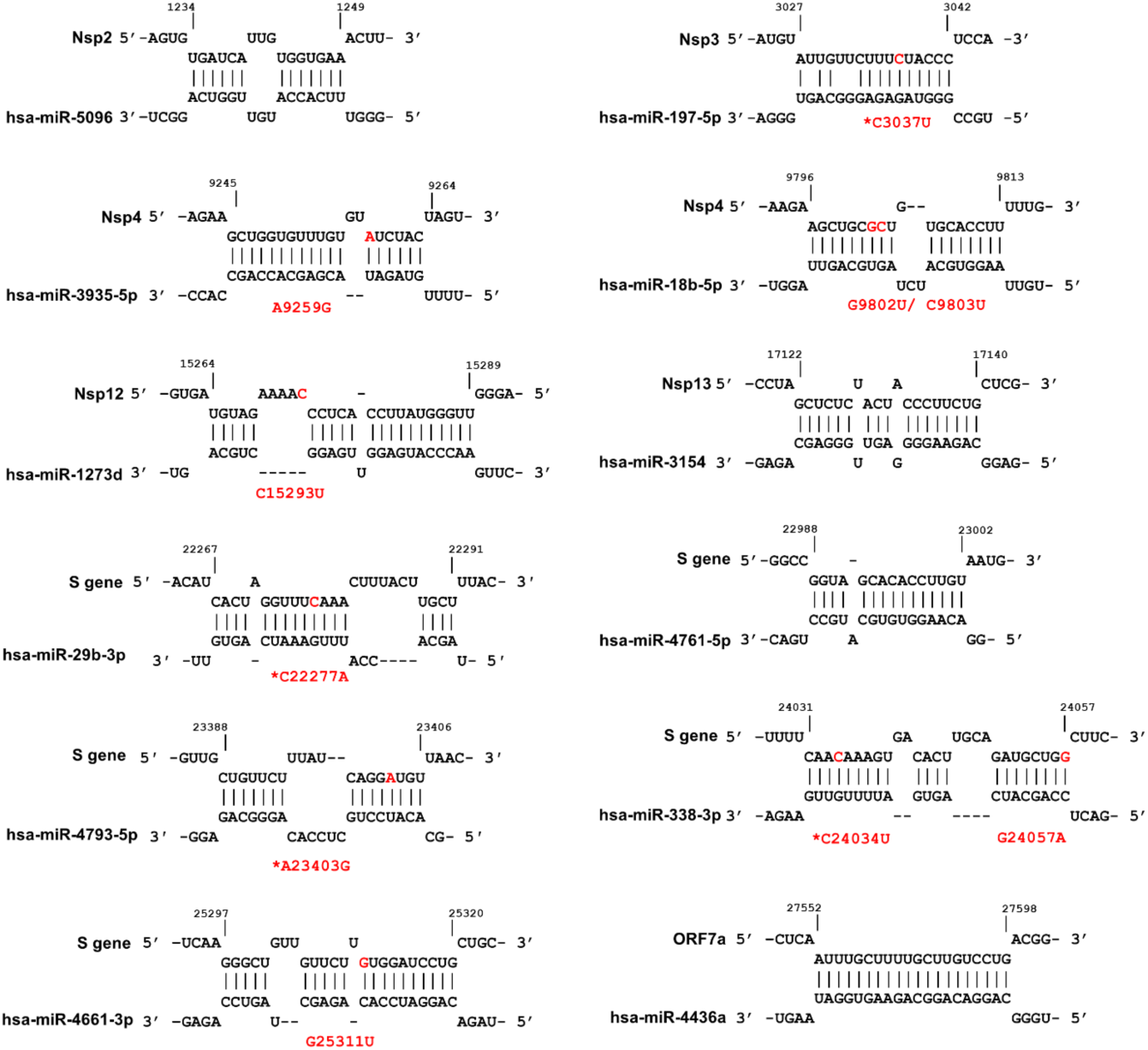
Prediction of host miRNAs binding sites within diffrenet regions of SARS-CoV-2 genome. The mutations that occur in miRNA binding sites are indicated in red, and the designations of the mutations are shown in red font and conserved mutations are indicated with an asterix. The figure was produced using IntaRNA tool.

MiR-197-5p is upregulated in patients with cardiovascular disease and its detection has been proposed as a biomarker [65-67]. In addition, it is well-known that patients with cardiovascular disease are overrepresented in symptomatic COVID-19 cohorts and have a higher mortality rate [68]. The C3037U conserved, but synonymous, mutation within Nsp3 sequence abolished the miR-197-5p target sequence (Figure 4). This mutation was introduced early January 2020 (Figure S1). This miRNA was previously reported to act in defence against hepatitis viruses, such as HBV, HCV, HAV and Enterovirus 71 [69-71].

Three mutations within Nsp4 result in loss of miR-3935 and miR-18b-5p target sequences, although they are not conserved mutations. Both miRNA are expressed in SARS-CoV-2 target cells (Figure 3B, S4 & S5). Nsp4 A9259G is in a sequence obtained from Vietnam (GISAID: EPI_ISL_416429), Nsp4 G9802U was reported in a Chinese patient [72] and Nsp4 C9803U was identified in the Netherlands (GISAID: EPI_ISL_422940) (Figure 4).

We identified five miRNAs with perfectly-matched complementary sequences within the S-gene: miR-29b-3p, miR-338-3p, miR-4661-3p, miR-4761-5p and miR-4793-5p (Figure 3). As shown in Figure 4, four of these sites were altered by recently identified mutations in the S-gene.

In particular, the miR-338-3p miRNA is expressed at high levels in SARS-CoV-2 target cells (Figure 3B, S9). The miR-338-3p target in the S-gene is removed by the C24034U conserved mutation identified early January in multiple countries (Figure S1). The miR-338-3p miRNA acts as tumor suppressor in liver, lung and gastric cancers [73-75]. The expression level of miR-338-3p declines during HBV infection [76,77] and miR-338-3p has a recognition site within Vaccinia virus genome [78]. Also, The G24057A mutation present in an sequence from Brazil (GISAID: EPI_ISL_429691), destroys the complementarity of the miR-338-3p binding site.

Of interest, two more conserved mutations C22277A (Q239K) and A23403G (D614G) occur in binding sites for miR-29b-3p and miR-4793-5p, respectively. Lastly, G25311U in a patient sample isolated India (MT396242.1) removed the miR-4661-3p binding site within the S gene (Figure 4).

In addition to the sites mentioned here, we identified an additional four host miRNAs with perfect complementarity within the receptor binding domain (RBD) region of S gene (Figure 5). These miRNA are not expressed by SARS-CoV-2 target cells (data not shown). However, because these sites exist within the critical ACE-2 targeting region, these miRNA target sequences may be relevant to miRNA-mediated virus attenuation technology. For example, viral replication can attenuated in a species-specific and tissue-specific manner by host miRNA machinery, which controls viral tropism, replication and pathogenesis [30-32].

**Figure 5.**
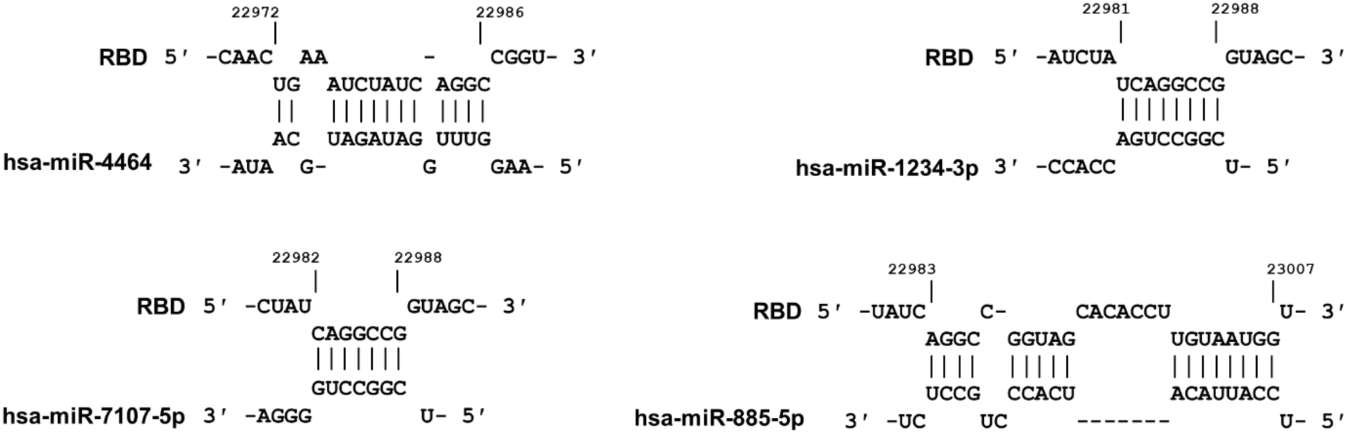
Identification of four miRNAs with ability to bind to the RBD within the S gene. These potential miRNAs can be used in miRNA-mediated attenuated virus technology.

### Possible impact of mutations on cryptic splice sites

Atypical cytoplasmic RNA splicing has been proposed to contribute to non-canonical viral transcripts, even for viruses that classically replicate in the cytoplasm [23-28]. Moreover, deep RNA sequencing has identified several previously unidentified SARS-CoV-2 viral RNAs that may be the result of non-canonical splicing events, or alternative transcriptional start sites [29]. We used RegRNA2 [37], HSF [38] and NIPU [39,40] tools to identify the putative splice sites and motifs within SARS-CoV-2 genome. Our computational prediction identified several 5′ donor and 3′ acceptor splice sites, as well as splice enhancer / inhibitor motifs [79] (Table 2). However, none of conserved mutations introduced, or deleted, any potential splice sites.

**Table 2.** Putative donor and acceptor splice sites and splice motifs in SARS-CoV-2 genome.

| Region | 5' Donor site | 3' Acceptor site | ESE | ESS | ISE | ISS |
| --- | --- | --- | --- | --- | --- | --- |
| 5' UTR | - | 1 | - | - | - | - |
| Nsp1 | 2 | - | 1 | 1 | 8 | - |
| Nsp2 | 1 | 1 | 3 | 2 | 6 | 1 |
| Nsp3 | 3 | 3 | 25 | 12 | 25 | - |
| Nsp4 | 4 | - | 6 | 1 | 6 | - |
| Nsp5 | 2 | 1 | 3 | 3 | 6 | - |
| Nsp6 | - | 3 | - | 1 | 5 | - |
| Nsp7 | - | - | 2 | 2 | - | - |
| Nsp8 | 1 | - | 5 | 1 | 2 | - |
| Nsp9 | 1 | - | 2 | - | 2 | - |
| Nsp10 | 1 | - | 2 | - | 1 | - |
| Nsp11 | - | - | - | - | 1 | - |
| Nsp12 | 1 | 5 | 9 | 7 | 10 | - |
| Nsp13 | 2 | 2 | 5 | 1 | 9 | - |
| **Nsp14** | 1 | 3 | 4 | 1 | 7 | - |
| **Nsp15** | - | - | 1 | 1 | - | - |
| **Nsp16** | - | 1 | 1 | 1 | - | - |
| **S** | - | 6 | 6 | 3 | 14 | 1 |
| **ORF3a** | - | 3 | 4 | 3 | 3 | - |
| **E** | 1 | 1 | - | - | 4 | - |
| **M** | - | 1 | 1 | 2 | 5 | - |
| **ORF6** | - | - | - | 1 | 1 | - |
| **ORF7a** | - | 1 | 1 | - | 3 | - |
| **ORF7b** | - | - | - | - | - | - |
| **ORF8** | 1 | - | 1 | - | 1 | - |
| **N** | 2 | 3 | 7 | 7 | 6 | - |
| **ORF10** | - | - | - | - | - | - |
| **3' UTR** | - | - | - | 1 | - | - |

ESE: exon splicing enhancer/ ESS: exon splicing silencer/ ISE: intron splicing enhancer/ ISS: intron splicing silencer.

## Discussion

At present there are nearly 200 mutations identified within global SARS-CoV-2 isolates. These mutations are mostly limited to point mutations, with little evidence for recombination events mediating the simultaneous transfer of multiple mutations. Although mutations may be due RdRp / Nsp12 infidelity, the predominance of C → U and G → A mutations is consistent with base-editing defence (e.g. APOBEC / ADAR) [57,80]. The Nsp14 exonuclease-based proof-reader is a critical counter-defence against host base-editor attack on the SARS-CoV-2 genome. It is also possible that the position of mutations within the genome could reflect accessibility of host base-editors to the SARS-CoV-2 genome upon uncoating, or during genome translation [57].

In our study, we filtered mutations to common / conserved events according to published sources [52]. There is little evidence that the existing mutations in SARS-CoV-2 have an impact on transmission, replication or viral load, but our study has flagged potential sites that could impact on viral fitness. It remains to be seen if these mutations undergo purifying selection in human populations over time. Carriage of SARS-CoV-2 mutations through the rapid expansion into naive populations throughout the world is most likely due to a neutral founder effects, rather than from fitness gains. For example, the high ratio of non-synonymous to synonymous mutations is close to the expected value for an emerging virus that has not undergone purifying selection [3].

One of the mutations (A23403G; D614G) in the Spike protein gene studied here defines the rapidly emerging G-clade present in one-third of global SARS-CoV-2 isolates. This mutation has been extensively studied by Korber *et al.* with respect to clinical outcomes and viral load [81]. The Korber et al. study showed that the A23403G did not enhance spike protein binding to ACE-2, however the mutation may be associated with higher viral load and poorer clinical outcomes. Our own analysis demonstrated that this mutation abolished a potential interaction with miR-4793-5p, a miRNA highly expressed in target cells including oesophagus, lung epithelium and small intestine. We propose that the A23403G mutation may represent an escape variant from the action of host miR-4793-5p.

Three mutations within Nsp4 were predicted to abolish miR-3935 and miR-18b-5p targeting. The expression of miR-3935 and miR-18b is altered upon viral infections [82-85]. The expression level of miR-3935 upregulates during H1N1, Crimean-Congo hemorrhagic fever virus, Coxsackievirus A16 and Enterovirus 71 infection [82,84,85]. On the other hand, miR-18b was reported to be downregulated during HBV and Ebola virus infections [83,86]. Similar to what observed for miR-197-5p, both miR-18b and miR-3935 are upregulated in patient with cardiovascular disease [87-89].

It is not yet clear if anti-viral miRNAs have evolved as host defense against viral infection, or are simply critical gene regulatory elements that assume an additional role for targeting viral transcripts – particularly when the human cellular defence machinery is confronted by an emerging zoonotic virus [9,13,14]. The possibility of including host miRNA binding sites into the genome of live-attenuated viruses offers a further checkpoint for the further attenuation of live vaccines, in a host-cell specific manner. For example, the identification of miRNA target sites in viral pathogens opens up opportunities for further study of viral host cell-tropism, or to create cell-specific or species-specific viral vaccines [30-32]. Finally, miRNA sites within the CDS of viral genes may be critical for ribosomal stalling, leading to the production of pioneer translation products (PTP). Enhanced production of PTP peptides may be critical for MHC-I loading for boosting the anti-viral CTL response [33-36].

## Conflict of interest

The authors declare that they have no conflicts of interest with the contents of this article.

## Author contribution

AH and AM conceived of the study and wrote the paper. AH performed the data analysis and produced the figures.

---

**Table S1.** Newly Identified common mutations with the counties of occurrence.

| **Mutation** | **Countries of occurrence** |
| --- | --- |
| **241** | Iran, Czech Republic, France, Germany, Greece, Israel, India, Nepal, Pakistan, Peru, South Africa, Spain, Taiwan and USA |
| **313** | Israel, Spain, India and USA |
| **1059** | Germany, Taiwan and USA |
| **14805** | Sir Lanka, Brazil, South Korea, Taiwan, Greece, Spain and USA |
| **17247** | Brazil, Sir Lanka and Taiwan |
| **25563** | France, Germany and USA |
| **28311** | Malaysia, South Korea, India, Germany and USA |
| **28688** | Iran, Taiwan, Turkey, Australia and Hong Kong |
| **29742** | Iran, Taiwan, Sir Lanka and Turkey |

**Figure S1.**
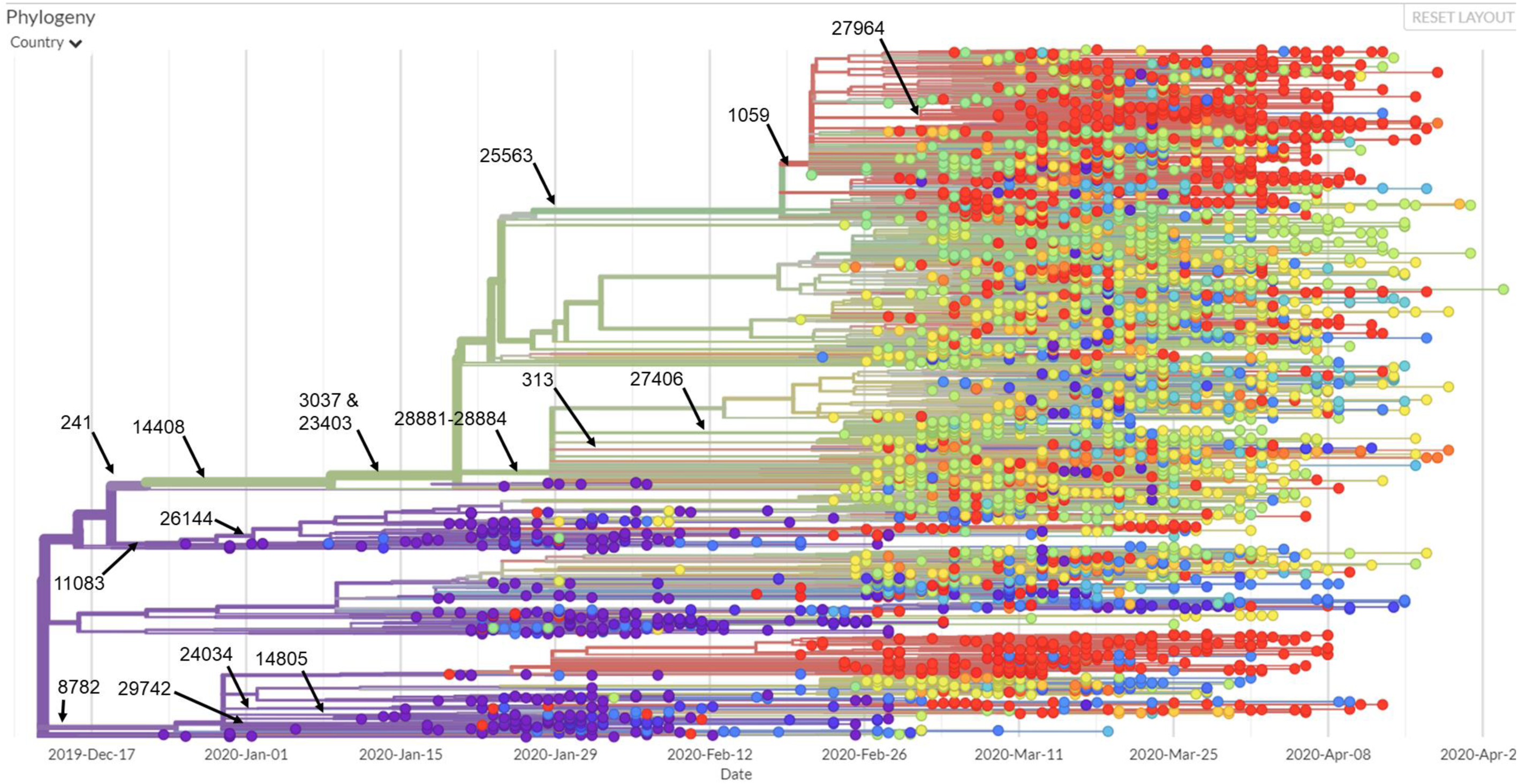
The position of conserve mutations in phylogeny graph obtained from Nextstrain database. Picture captured on May 1st, 11: 21 AM.

**Figure S2.**
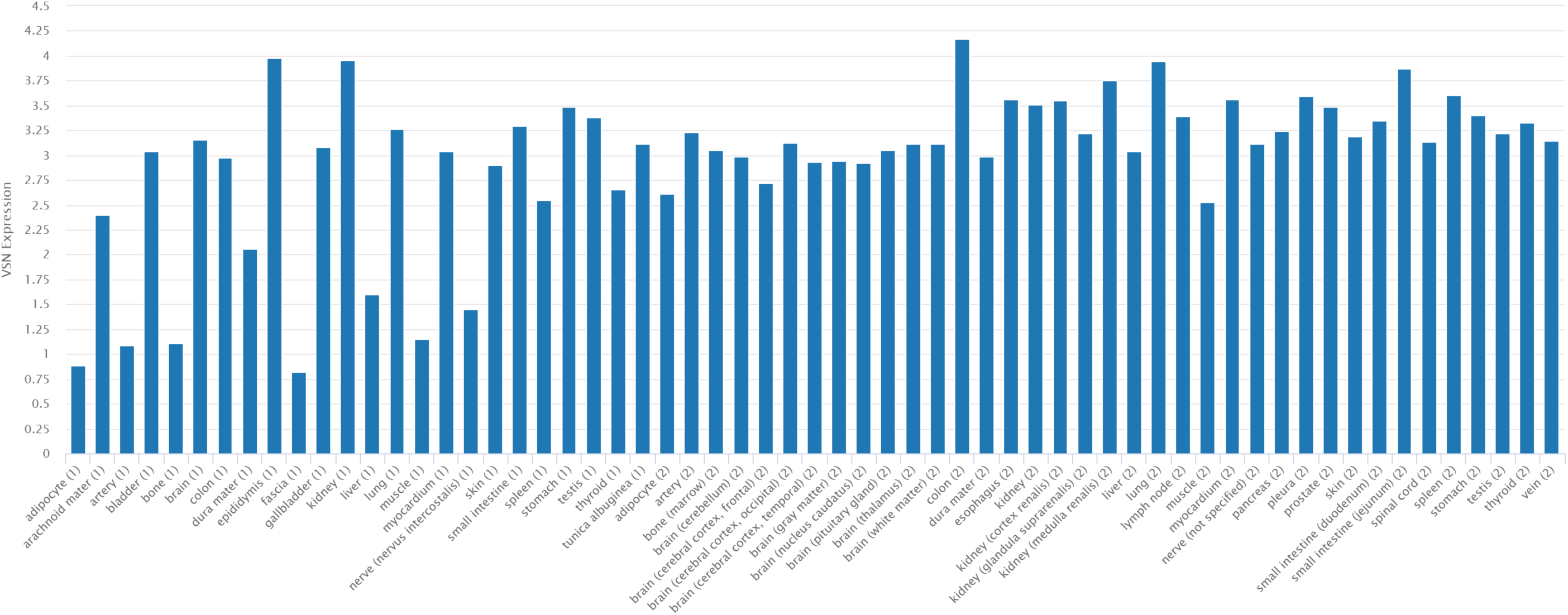
The expression of miR-5096 in different human tissue obtained from TissueAtlas data base. The graph represents expression level in two different samples.

**Figure S3.**
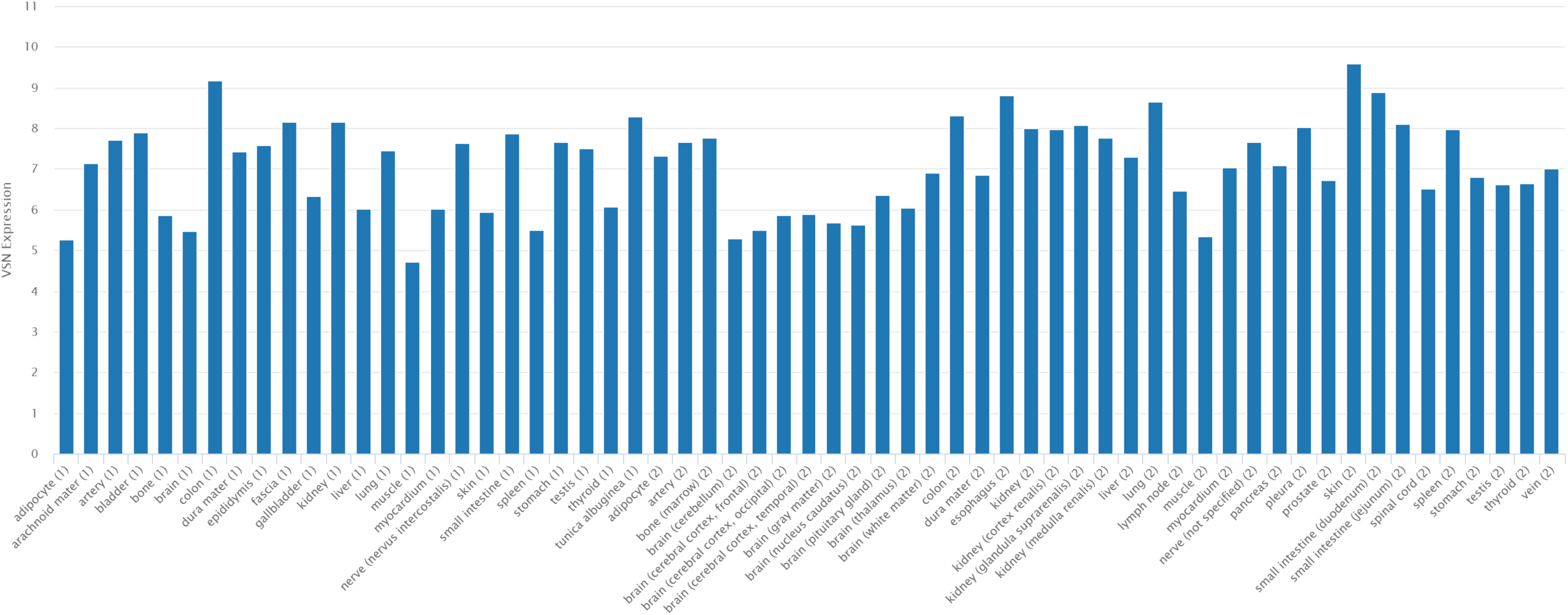
The expression of miR-197-5p in different human tissue obtained from TissueAtlas data base. The graph represents expression level in two different samples.

**Figure S4.**
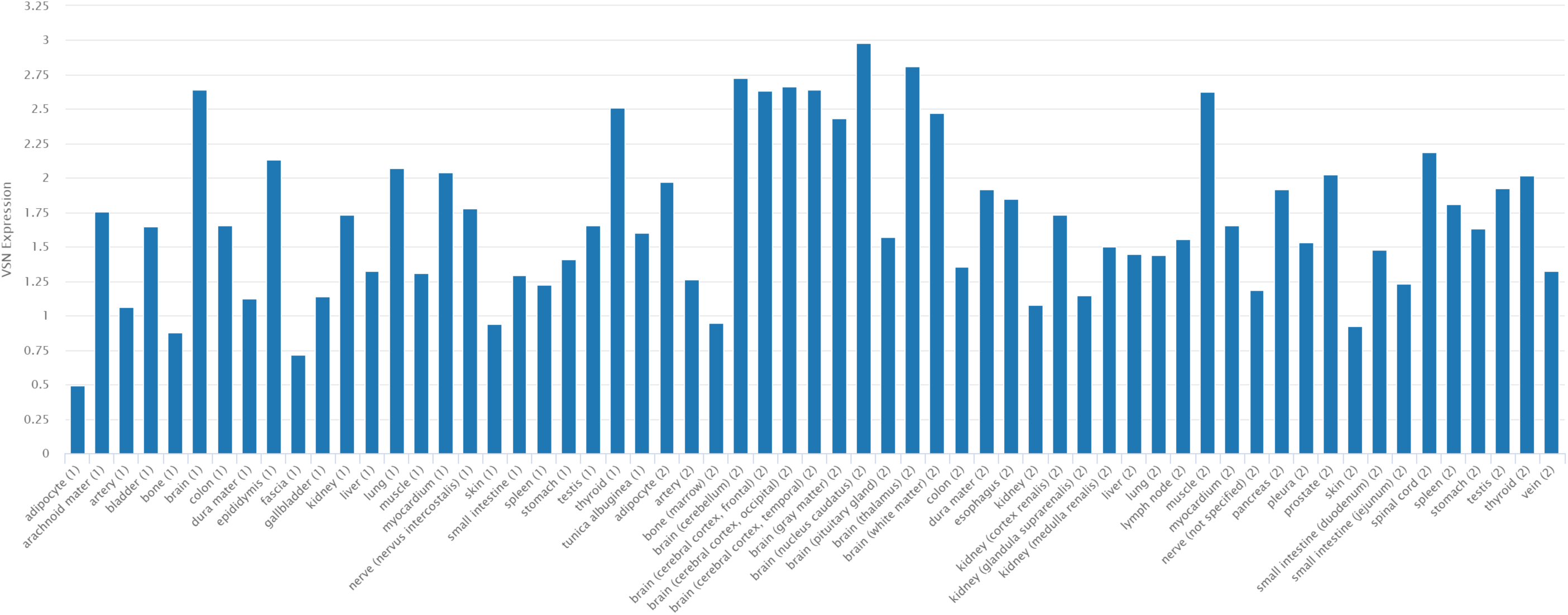
The expression of miR-3935 in different human tissue obtained from TissueAtlas data base. The graph represents expression level in two different samples.

**Figure S5.**
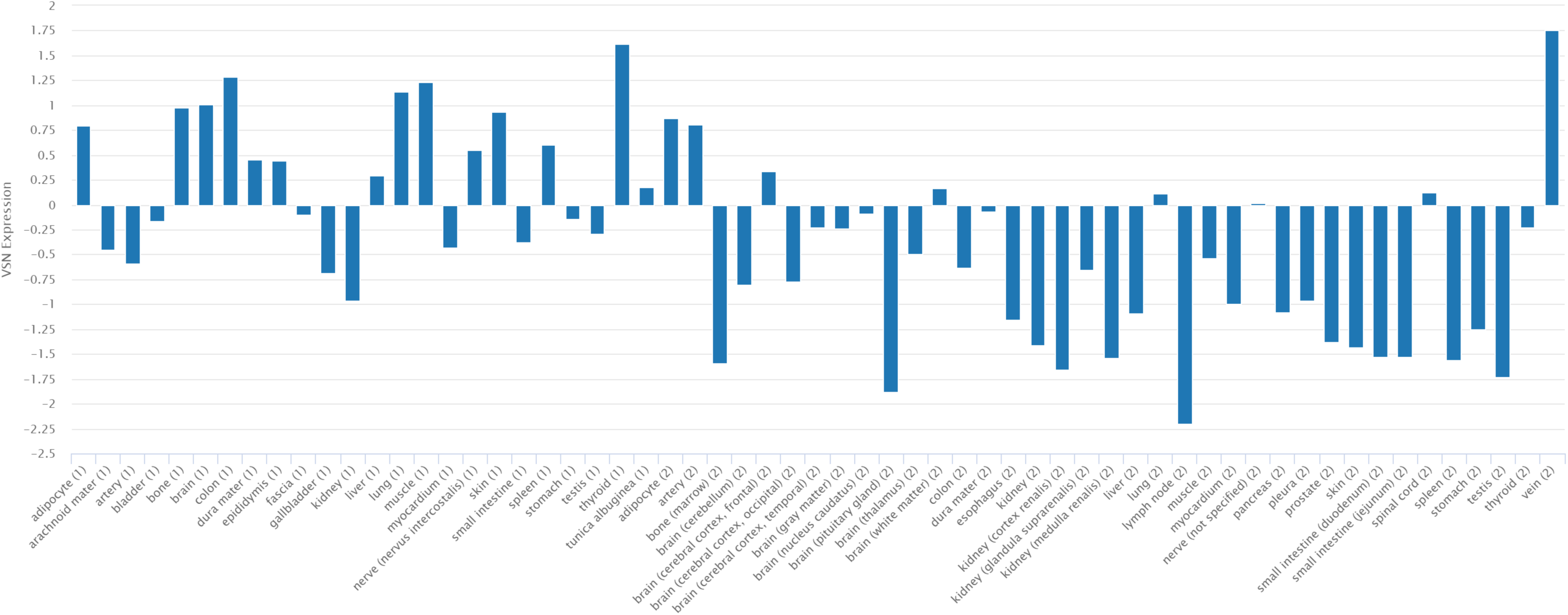
The expression of miR-18b-5p in different human tissue obtained from TissueAtlas data base. The graph represents expression level in two different samples.

**Figure S6.**
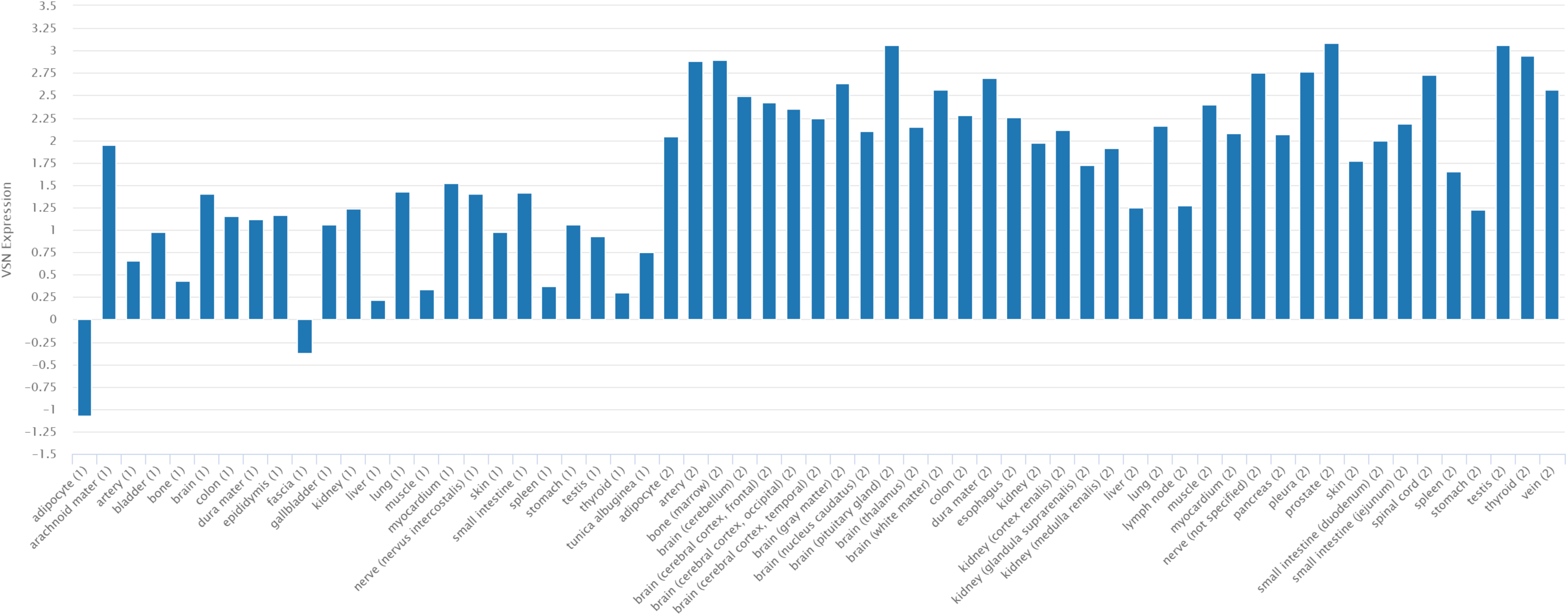
The expression of miR-1273d in different human tissue obtained from TissueAtlas data base. The graph represents expression level in two different samples.

**Figure S7.**
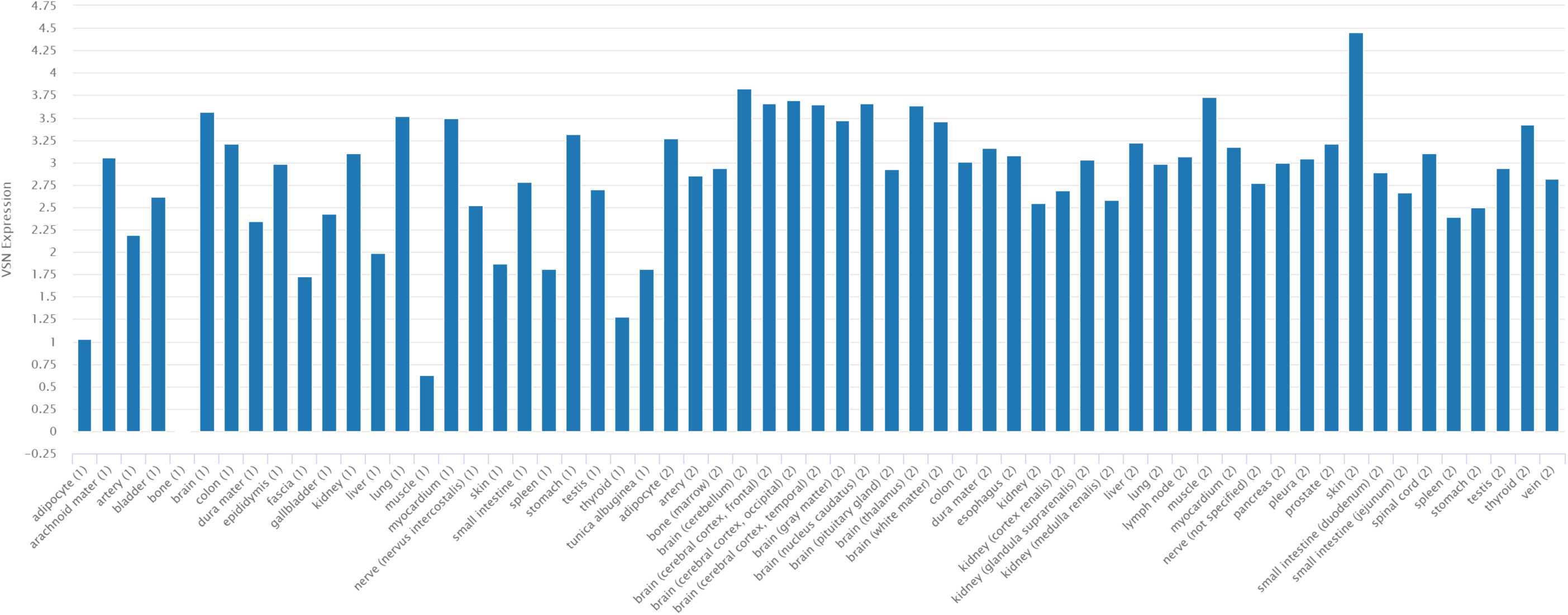
The expression of miR-3154 in different human tissue obtained from TissueAtlas data base. The graph represents expression level in two different samples.

**Figure S8.**
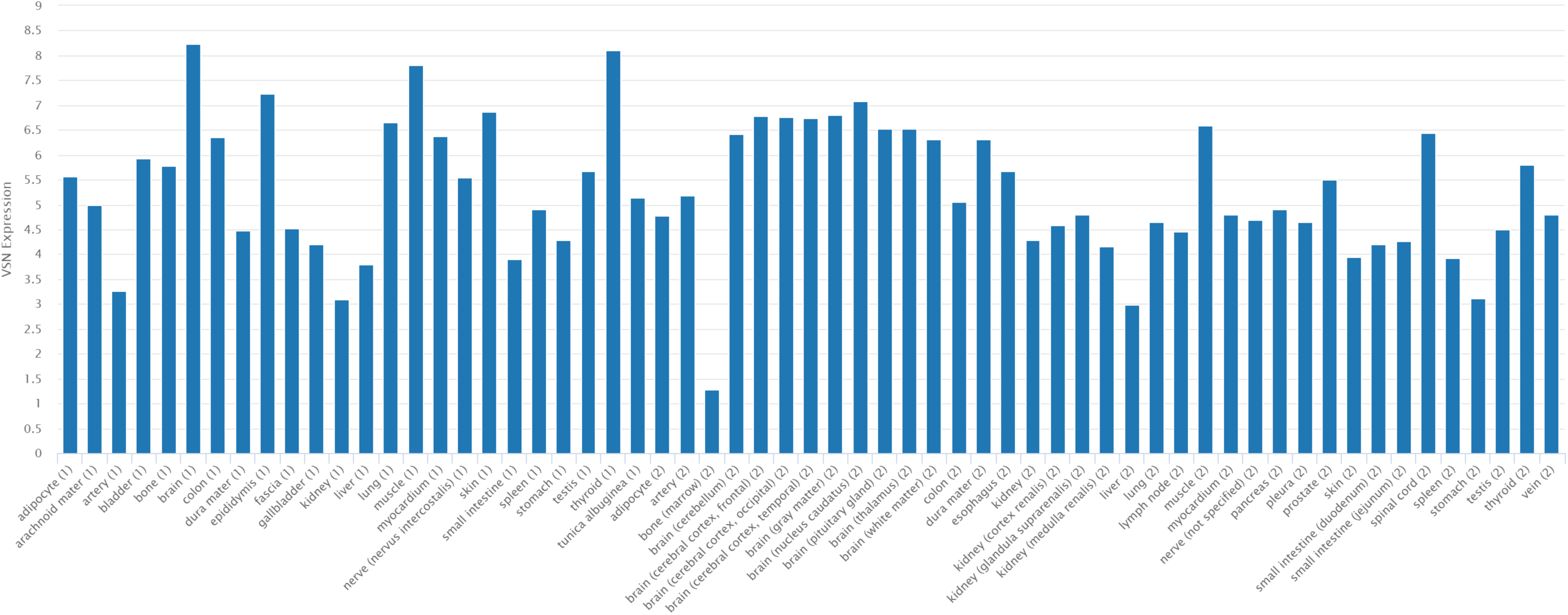
The expression of miR-29b-3p in different human tissue obtained from TissueAtlas data base. The graph represents expression level in two different samples.

**Figure S9.**
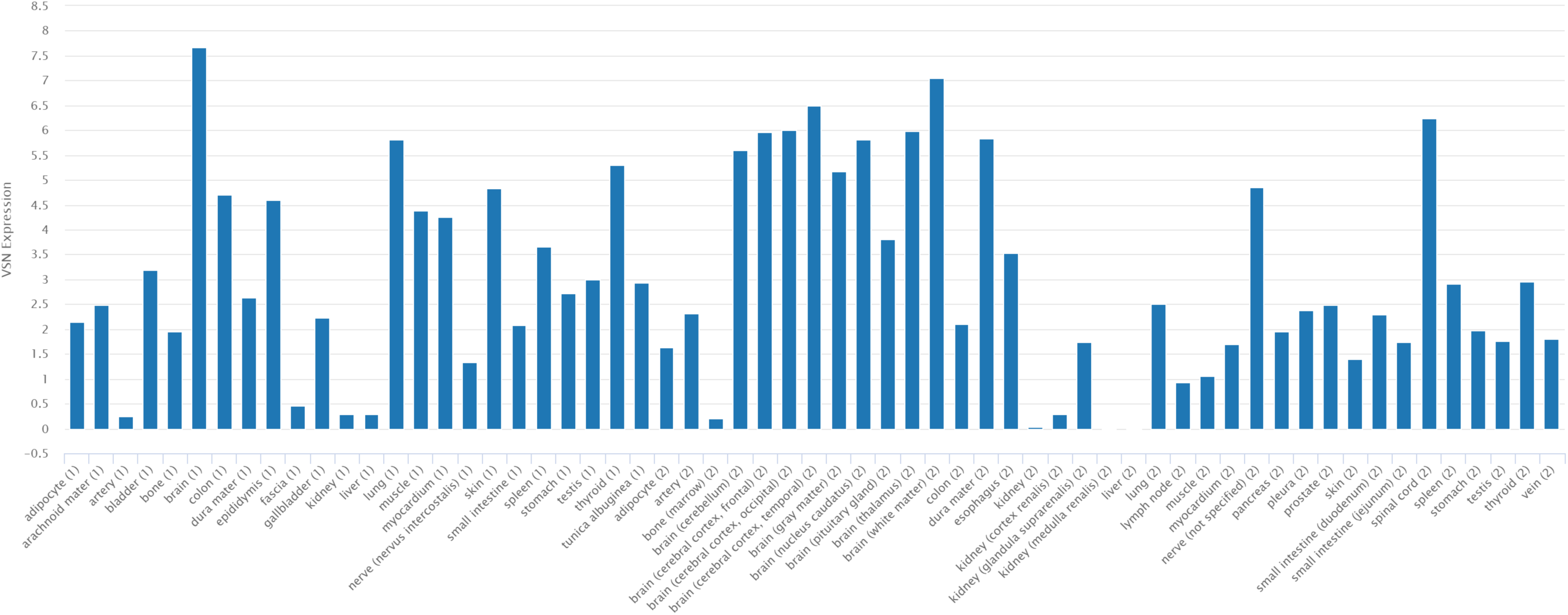
The expression of miR-338-3p in different human tissue obtained from TissueAtlas data base. The graph represents expression level in two different samples.

**Figure S10.**
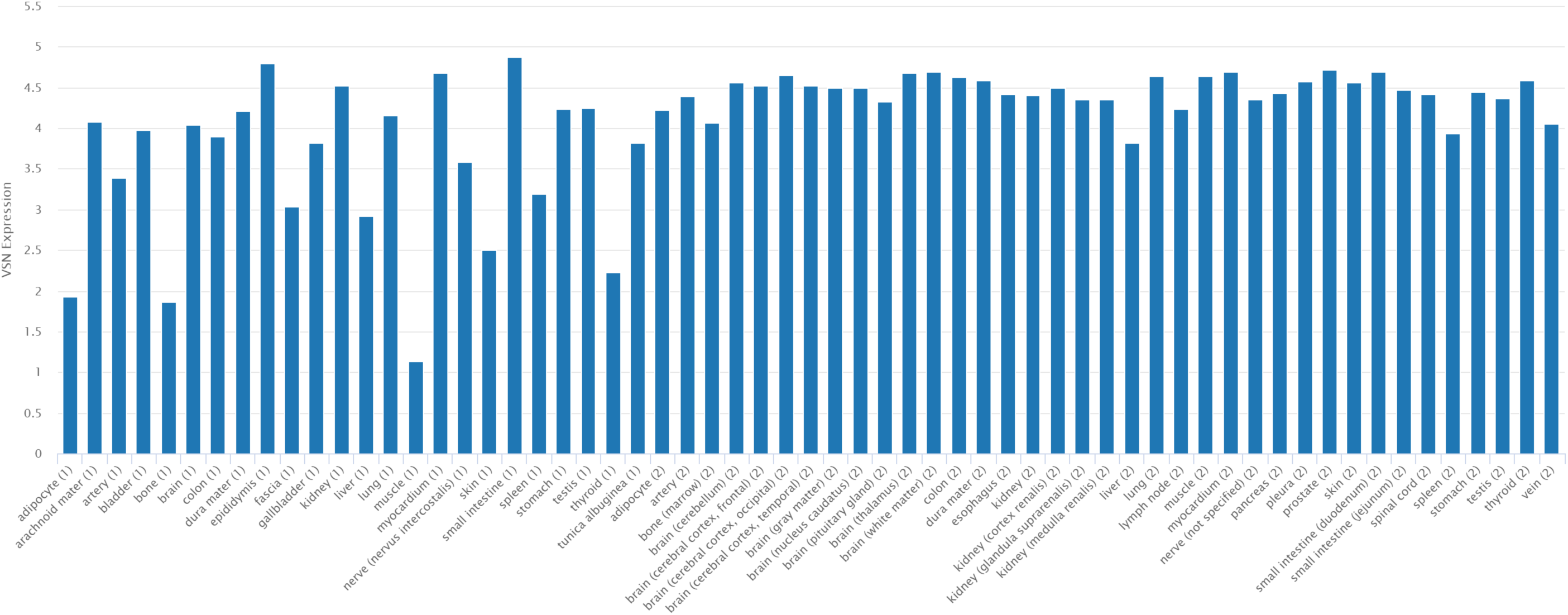
The expression of miR-4793-5p in different human tissue obtained from TissueAtlas data base. The graph represents expression level in two different samples.

**Figure S11.**
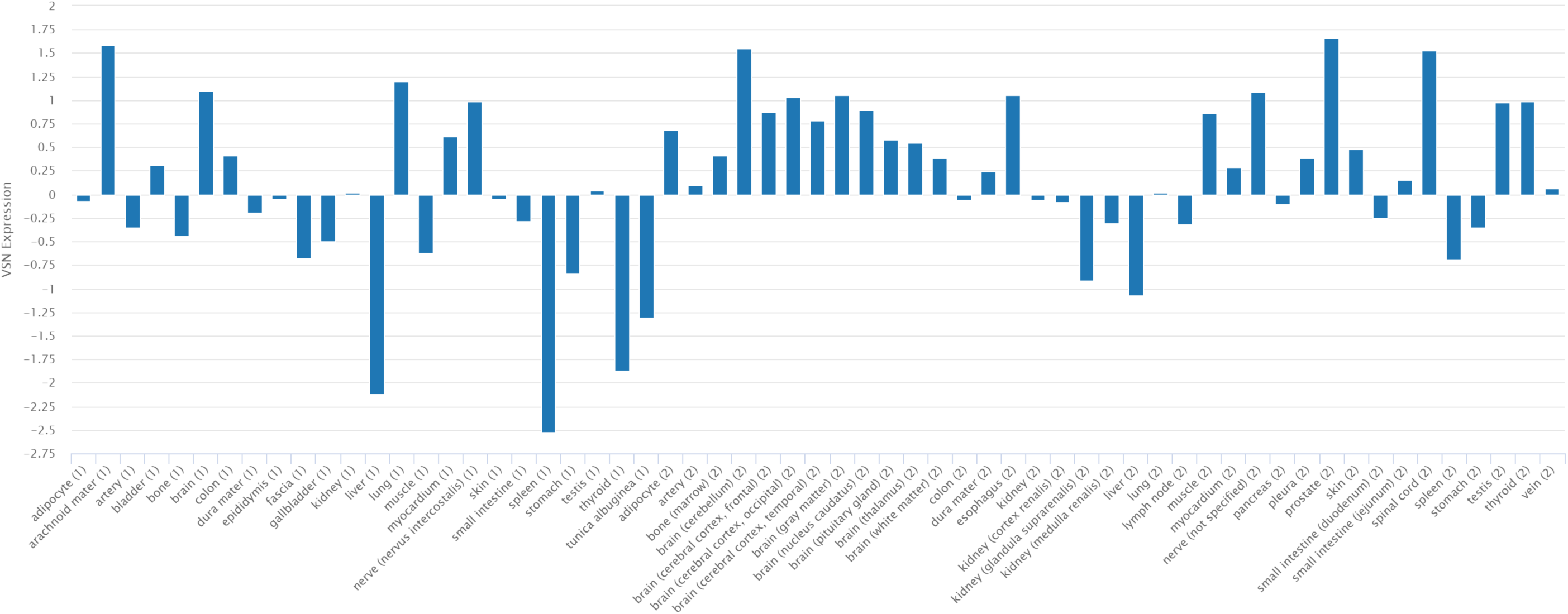
The expression of miR-4436a in different human tissue obtained from TissueAtlas data base. The graph represents expression level in two different samples.

## Notes

### Competing Interest Statement

The authors have declared no competing interest.

